# Genetic Regulators of Sputum Mucin Concentration and Their Associations with COPD Phenotypes

**DOI:** 10.1101/2022.09.28.509838

**Authors:** Eric Van Buren, Giorgia Radicioni, Sarah Lester, Wanda K. O’Neal, Hong Dang, Silva Kasela, Suresh Garudadri, Jeffrey L. Curtis, Meilan Han, Jerry A. Krishnan, Emily S. Wan, Edwin K. Silverman, Annette Hastie, Victor E. Ortega, Tuuli Lappalainen, Stephanie A. Christenson, Yun Li, Michael H. Cho, Mehmet Kesimer, Samir N. P. Kelada

## Abstract

Hyper-secretion and/or hyper-concentration of mucus is a defining feature of multiple obstructive lung diseases, including chronic obstructive pulmonary disease (COPD). Mucus itself is composed of a mixture of water, ions, salt and proteins, of which the gel-forming mucins, MUC5AC and MUC5B, are the most abundant. Recent studies have linked the concentrations of these proteins in sputum to COPD phenotypes, including chronic bronchitis (CB) and acute exacerbations (AE). We sought to determine whether common genetic variants influence sputum mucin concentrations and whether these variants are also associated with COPD phenotypes, specifically CB and AE. We performed a GWAS to identify quantitative trait loci for sputum mucin protein concentration (pQTL) in the Sub-Populations and InteRmediate Outcome Measures in COPD Study (SPIROMICS, n=708 for total mucin, n=215 for MUC5AC, MUC5B). Subsequently, we tested for associations of mucin pQTL with CB and AE using regression modeling (n=822-1300). Replication analysis was conducted using data from COPDGene (n =5740) and by examining results from the UK Biobank. We identified one genome-wide significant pQTL for MUC5AC (rs75401036) and two for MUC5B (rs140324259, rs10001928). The strongest association for MUC5B, with rs140324259 on chromosome 11, explained 14% of variation in sputum MUC5B. Despite being associated with lower MUC5B, the C allele of rs140324259 conferred increased risk of CB (odds ratio (OR) = 1.42; 95% confidence interval (CI): 1.10-1.80) as well as AE ascertained over three years of follow up (OR=1.41; 95% CI: 1.02-1.94). Associations between rs140324259 and CB or AE did not replicate in COPDGene. However, in the UK Biobank, rs140324259 was associated with phenotypes that define CB, namely chronic mucus production and cough, again with the C allele conferring increased risk. We conclude that sputum MUC5AC and MUC5B concentrations are associated with common genetic variants, and the top locus for MUC5B may influence COPD phenotypes, in particular CB.

**Author Summary:** Chronic obstructive pulmonary disease (COPD) is characterized by presence of emphysema and/or chronic bronchitis. Excessive mucus production is a defining phenotype of chronic bronchitis, and is associated with several important features of COPD, including exacerbations and loss of lung function. Recent studies have demonstrated that the amount of mucus produced in COPD patients is an important marker of disease state. We investigated whether common genetic variants are associated with the concentration of two key proteins in mucus, MUC5AC and MUC5B, and whether the variants we identified are also associated with COPD outcomes. We identified multiple genetic variants that were associated with MUC5AC or MUC5B concentration. The strongest association we detected, for MUC5B on chromosome 11, was also associated with features of COPD, including chronic bronchitis and acute exacerbations, in one COPD study population but not another. Results from a much larger study, the UK Biobank, indicate that this variant is associated with chronic mucus production and chronic cough, which are key features of chronic bronchitis. Thus, we conclude that the concentration of key proteins in mucus are influenced by genetic variation, and that a variant on chromosome 11 that affects MUC5B may in turn alter COPD outcomes.

## INTRODUCTION

Chronic obstructive pulmonary disease (COPD) is a smoking-related disease that affects more than 200 million people and is the fourth leading cause of death worldwide [1,2]. The disease is characterized by the presence of emphysema and/or chronic bronchitis (CB). Chronic mucus hyper-secretion is a defining phenotype of CB and is associated with airway obstruction due to mucus plugs [3], acute exacerbations (AE) [4], and accelerated loss of lung function over time [5,6].

Mucus itself is composed of a mixture of water, ions, salt and proteins, and mucin glycoproteins, most prominently the gel-forming mucins, MUC5AC and MUC5B. Their concentrations and biochemical properties (e.g., size and oxidation state) largely determine the viscoelastic properties of mucus in health and disease [7]. Recent studies emanating from the Sub-Populations and InteRmediate Outcome Measures in COPD Study (SPIROMICS) have provided compelling evidence that the concentration of mucin proteins in induced sputum is an important biomarker in COPD [8,9]. Kesimer et al. showed that mucin concentrations (total, MUC5AC and/or MUC5B) was associated with smoking history, phlegm production, CB, risk of AE, and disease severity [8]. A subsequent study showed that the concentrations of MUC5AC was associated with disease initiation and progression [9].

Here, we report our findings of a genome-wide search for common genetic variants associated with variation in sputum mucin protein concentration, that is, mucin protein quantitative trait loci (pQTL). Previous studies have identified genetic variants associated with mucin gene expression (eQTL) that are located within or near *MUC5AC* [10–12] (in asthma) and *MUC5B* (in idiopathic pulmonary fibrosis, IPF [13]). That said, we conducted a genome-wide search for mucin pQTL because our prior work in a mouse model system indicated distal (or trans) pQTL for mucins were possible and perhaps even likely [14]. We leveraged quantitative mass spectrometry-based measurements of samples from SPIROMICS that were generated previously [8,9], to identify main effect pQTL and pQTL that result from genotype × smoking interactions. Subsequently, we tested whether the pQTL we identified were associated with COPD outcomes, namely CB and AE, in SPIROMICS, followed by replication analysis in COPDGene and the UK Biobank.

## RESULTS

### GWAS for Mucin pQTL

We conducted a GWAS of total and specific (MUC5AC and MUC5B) mucin concentrations in sputum to identify novel regulators of these biomarkers in SPIROMICS. Descriptive statistics of study participants are provided in Table 1. The mucin phenotype data represent a subset of subjects described in two previous studies [8,9], and comprise a subset of participants in SPIROMICS (Figure S1). In this sample, there was a clear effect of smoking history on total mucin concentration (Figure S2), but among COPD cases, there was not a linear or monotonically increasing relationship between total mucin concentration and disease severity (as reflected by Global Initiative for Chronic Obstructive Lung Disease (GOLD) stage). Similar patterns were observed for MUC5AC and MUC5B (Figure S2).

**Table 1.** Descriptive Statistics of SPIROMICS Participants in Mucin GWAS*.

|  | **Non-smoking Controls** |  | **At-risk** |  | **GOLD 1** |  | **GOLD 2** |  | **GOLD 3** |  |
| --- | --- | --- | --- | --- | --- | --- | --- | --- | --- | --- |
| Study Population | Total Mucin | MUC5AC, MUC5B | Total Mucin | MUC5AC, MUC5B | Total Mucin | MUC5AC, MUC5B | Total Mucin | MUC5AC, MUC5B | Total Mucin | MUC5AC, MUC5B |
| N | 50 | 25 | 219 | 46 | 130 | 40 | 241 | 56 | 67 | 48 |
| Age, mean (range) | 57.9 (40-80) | 61.3 (42-80) | 60.4 (40-79) | 63.3 (40-79) | 66.7 (45-80) | 65.8 (49-79) | 65.0 (42-80) | 64.9 (48-80) | 66.5 (48-80) | 67.1 (52-80) |
| Males, n (%) | 26 (51.0) | 13 (52.0) | 117 (53.2) | 21 (45.7) | 91 (70.0) | 31 (77.5) | 142 (58.9) | 35 (62.5) | 35 (52.2) | 21 (43.8) |
| European Ancestry, n (%) | 36 (72.0) | 25 (100) | 155 (70.8) | 46 (100) | 112 (86.2) | 40 (100) | 214 (88.8) | 56 (100) | 59 (88.1) | 48 (100) |
| African Ancestry, n (%) | 14 (28.0) | NA<sup>†</sup> | 64 (29.2) | NA<sup>‡</sup> | 18 (13.9) | NA<sup>‡</sup> | 27 (11.2) | NA<sup>‡</sup> | 8 (11.9) | NA<sup>‡</sup> |
| Chronic Bronchitis<sup>†</sup>, n (%) | 3 (6.0) | 2 (8.0) | 89 (40.6) | 15 (32.6) | 52 (40.0) | 19 (47.5) | 134 (55.4) | 35 (62.5) | 29 (43.2) | 20 (41.7) |
| Current smoker, n (%) | 0 (0) | 0 (0) | 113 (51.6) | 21 (45.7) | 45 (34.6) | 17 (42.5) | 114 (47.1) | 26 (46.4) | 18 (26.9) | 11 (22.9) |
| Former smoker, n (%) | 0 (0) | 0 | 106 (48.4) | 25 (54.4) | 85 (65.4) | 23 (57.5) | 127 (52.5) | 30 (53.6) | 49 (73.3) | 37 (77.1) |
| Smoking, pack-years (range) | 0 | 0 (0) | 43.0 (20-150) | 44.0 (20-100) | 51.9 (20-160) | 54.7 (20-117) | 57.1 (20-270) | 55.8 (20-147) | 51.8 (20-126) | 50.3 (21-126) |

\*One subject with a GOLD stage of 4 was included in the total mucin analysis but is not shown here.
<sup>†</sup> Based on St. George's Respiratory Questionnaire
<sup>‡</sup>Due to limited sample size of SPIROMICS participants of African ancestry with MUC5AC/MUC5B data, only data on European ancestry subjects were used in these analyses.

We did not detect any genome-wide significant loci (p<5.0×10^−8^) associated with total mucin concentration based on data from SPIROMICS participants of European ancestry (EA, N=576) or African ancestry (AA, N=132) participants (Figures S3 and S4), nor in combined analysis of EA and AA subjects (not shown). Testing for joint effects of SNP and SNP × smoking (pack-years) interactions did not reveal any loci associated with total mucin concentration either. In contrast, despite relatively small sample size (N=215 EA subjects), we identified three genome-wide significant pQTL for MUC5AC or MUC5B (Figure 1, Table S1, and Figures S5 and S6). In addition to the pQTL for MUC5AC on chromosome (Chr) 7 (rs75401036), we identified one highly suggestive locus on Chr 2 (rs16866419, p=7.2×10^−8^), thus both MUC5AC pQTL are located on chromosomes other than Chr 11 where *MUC5AC* and *MUC5B* are located (i.e., act in trans). We note that restricting our analysis to variants located in/near MUC5AC, i.e., with a reduced multiple testing correction, did not reveal any local pQTL for MUC5AC, and this includes testing variants previously associated with *MUC5AC* gene expression [10–12] including rs12788104, rs11602802, rs11603634, rs1292198170, and rs1132436. Overall, SNP-based heritability estimates for MUC5AC (*h*^2^_SNP_=0.712; S.E. = 1.322) and MUC5B (*h*^2^_SNP_ = 0.608, S.E.=1.475) were high but imprecise, which is not surprising given the relatively small sample size.

**Figure 1.**
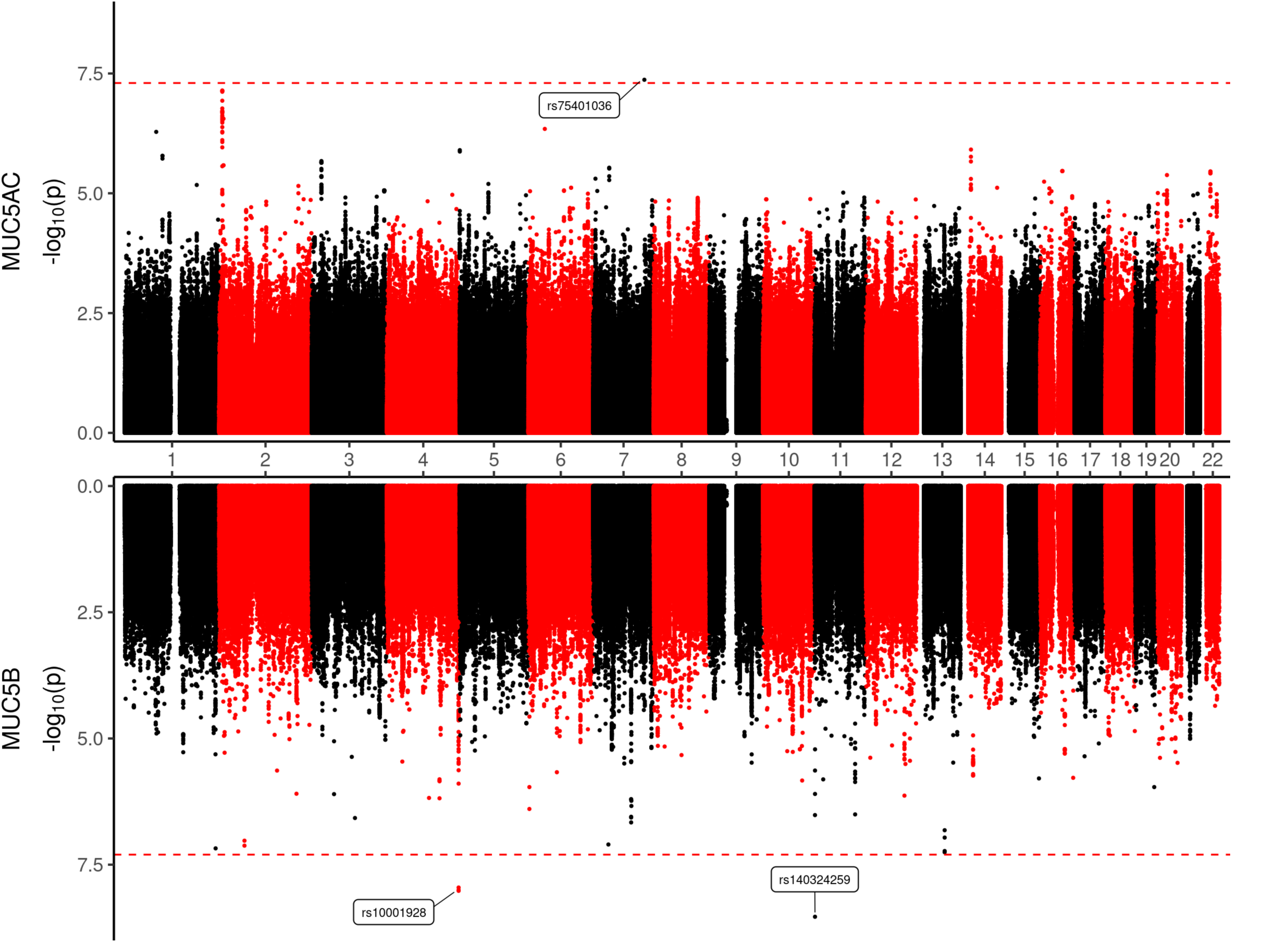
Distal and local pQTL for sputum MUC5AC and MUC5B. Results of association analysis using sputum mucin concentration data from 215 EA SPIROMICS participants are shown. Dashed red line denotes genome-wide significance threshold.

For MUC5B, one local pQTL was detected on Chr 11 (rs140324259), and one distal pQTL was located on Chr 4 (rs10001928). One additional MUC5B pQTL (rs6043852), located in the intron of *KIF16B* on chromosome 20 was detected by testing for the joint effects of SNP + SNP × smoking interactions (joint test p-value = 1.3×10^−9^, Figure 2A). Further analysis revealed that the interaction itself contributed substantially to the joint effect (p_interaction_ = 1.1×10^−7^), such that rs6043852 was associated with MUC5B concentration only in subjects that are not current smokers (Figure 2B). Given the relatively low minor allele frequency of rs6043852 (3%), the number of subjects harboring genotypes with the minor allele (A) and of contrasting smoking status was not large (n=6 A allele carriers in current smokers and n=6 in the combined never plus former smoker group). Hence this SNP × smoking interaction pQTL must be interpreted with caution.

**Figure 2.**
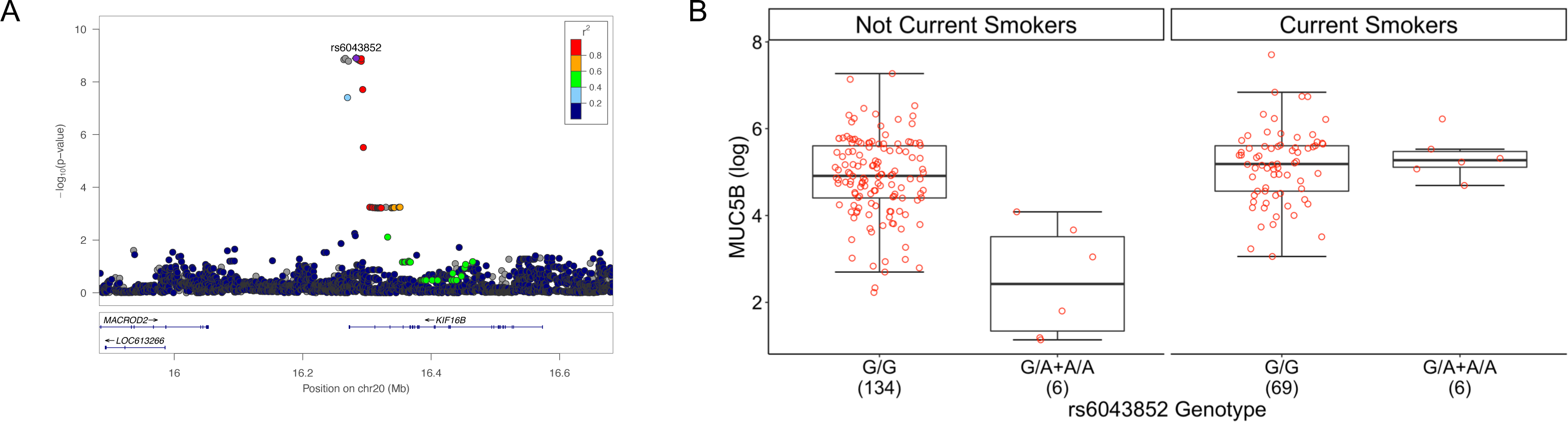
A Genotype x Smoking interaction locus (rs6043852) for sputum MUC5B on Chromosome 20. **A.** Locus zoom plot for the genotype x smoking locus (rs6043852). **B**. MUC5B concentration as a function of both rs6043852 genotype and current smoking status. The not current smoker category includes never smokers and former smokers. Note that while we plot carriers of the minor allele here as one group, the regression model for MUC5B used genotype dosages.

The strongest pQTL we identified was for MUC5B on Chr 11. The lead variant, rs140324259, is located approximately 100 kb upstream of *MUC5B*, in between *MUC2* and *MUC5AC* (Figure 3A). A second variant located in intron 6 of *MUC5AC*, rs28668859, was also associated with MUC5B concentration; conditional analysis revealed this this signal was partially dependent on linkage disequilibrium (LD, R^2^=0.20) with rs140324259 (conditional p-value = 1.6×10^−3^). Neither rs140324259 nor rs28668859 are in LD (R^2^=0.02 and 0.01, respectively) with the *MUC5B* promoter variant rs35705950 that is a well-known *MUC5B* eQTL and is associated with IPF [13]. After adjusting for covariates, rs140324259 genotype explained ~14% of variation in sputum MUC5B, and each minor allele (C) contributed a ~2.3 pmol/ml unit decrease in MUC5B (Figure 3B), an effect size that is greater than the effect of current smoking status (yes vs. no, ~1.4 pmol/ml). Adjusting for disease severity (using GOLD stage) did not materially change these results.

**Figure 3.**
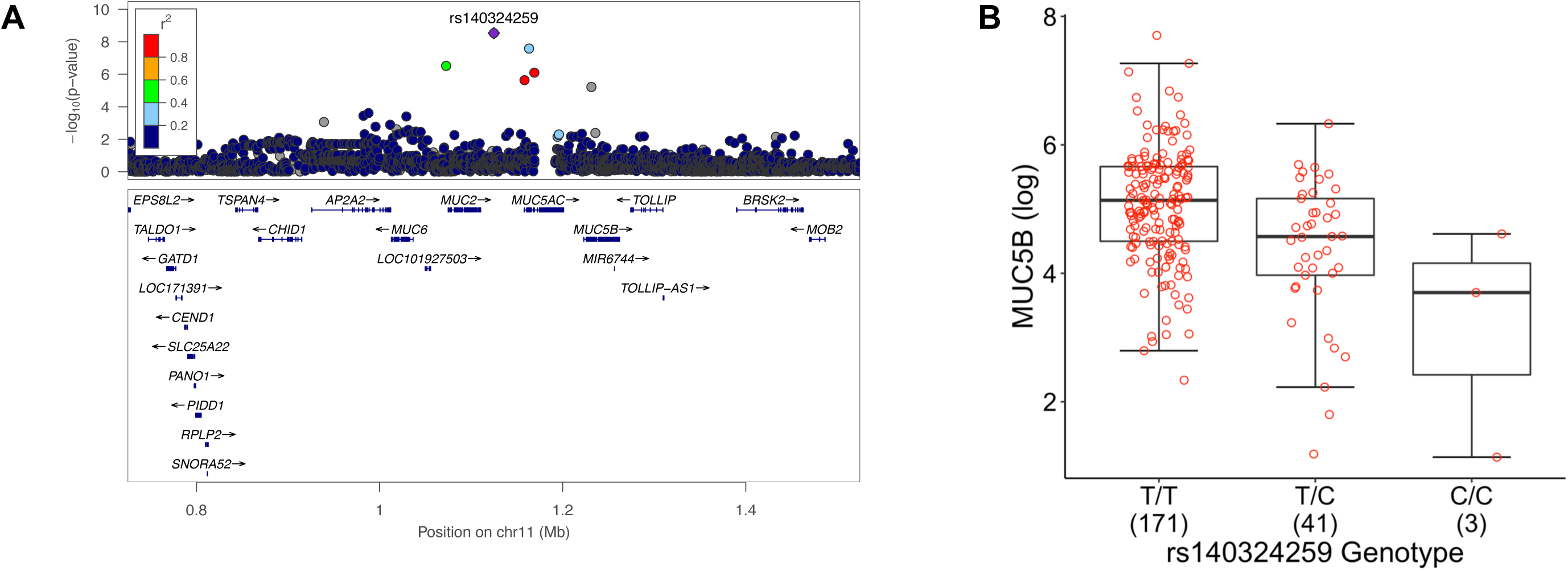
The Chromosome 11 MUC5B pQTL. **A**. Regional view of association test results. Four genes were omitted due to small size. The lead variant, rs140324259, is approximately 100 kb upstream of *MUC5B*. **B**. Effect of rs140324259 genotype on sputum MUC5B. Numbers in parentheses on x-axis denote sample size per genotype. Each C allele yields a 0.8 log (ln) unit decreased in MUC5B, corresponding to 2.3 picomol/ml drop in MUC5B concentration.

We asked whether lead MUC5B pQTL, rs140324259, was associated with *MUC5B* gene expression in the SPIROMICS participants or other studies. In a subset of SPIROMICS participants (n=144) for whom airway brush RNA-seq data exist [15], we used a tagSNP for rs140324259, namely rs55680540 (LD R^2^=0.72 in entire SPIROMICS population), but found no correlation between genotype and *MUC5B* expression (Figure S7A). No other variants in the region were significantly associated with *MUC5B* expression (FDR = 0.497, Figure S7B). Additionally, rs140324259 was also not associated with *MUC5B* expression in the nasal epithelium of subjects with cystic fibrosis [16] or asthma [12], nor was it reported as an eQTL in any tissue in the GTEx dataset [17], including homogenized lung tissue.

Given that power for eQTL detection could be an issue underlying the negative eQTL association results, we asked whether rs140324259 or four variants in LD (rs55680540, rs28668859, rs11604917, and rs76498418) could potentially affect gene expression by altering transcription factor binding or chromatin state using Haploreg [18]. As shown in Table S2, rs140324259, rs11604917, and rs55680540 are predicted to alter binding of transcription factors, and there is some evidence that rs11604917 and rs55680540 alter chromatin state in relevant cell types or tissues (Table S3). Perhaps most notably, rs11604917 lies in an enhancer region in multiple cell types and tissues and is predicted to alter binding of the transcription factor RBP-J. The alternate allele (C) of rs11604917 disrupts the consensus sequence at the first position of an almost invariant motif (Figure S8). Given that RBP-J is part of the Notch signaling pathway that determines ciliated vs. secretory cell fate in murine airways [19], this finding potentially merits further investigation.

### Association of MUC5B pQTL with COPD Phenotypes

Subsequently, we tested whether rs140324259 was related to clinically-relevant COPD phenotypes, namely CB and AE. In the subset of SPIROMICS participants with complete phenotype, genotype, sputum MUC5B, and clinical outcomes data (n=141), we found that rs140324259 was not associated with CB at baseline/enrollment (p=0.25, Table S4). rs140324259 was not associated with AE in the year prior to enrollment (p=0.14, Figure 4A) unless we accounted for sputum MUC5B concentration (p=0.02, Figure 4B, and Table S5). Surprisingly, in this analysis, we found that while MUC5B concentration was positively associated with AE (β_1_=0.45, p=0.01, Figure 4B), the effect of rs140324259 genotype (β_2_=0.74, p=0.02) was opposite our expectation based on pQTL analysis. That is, the C allele of rs140324259, which was associated with lower MUC5B (γ =−0.77, p=2.6 ×10^−6^) and therefore would be expected to confer decreased risk of AE, was associated with increased risk of AE compared to the T allele.

**Figure 4.**
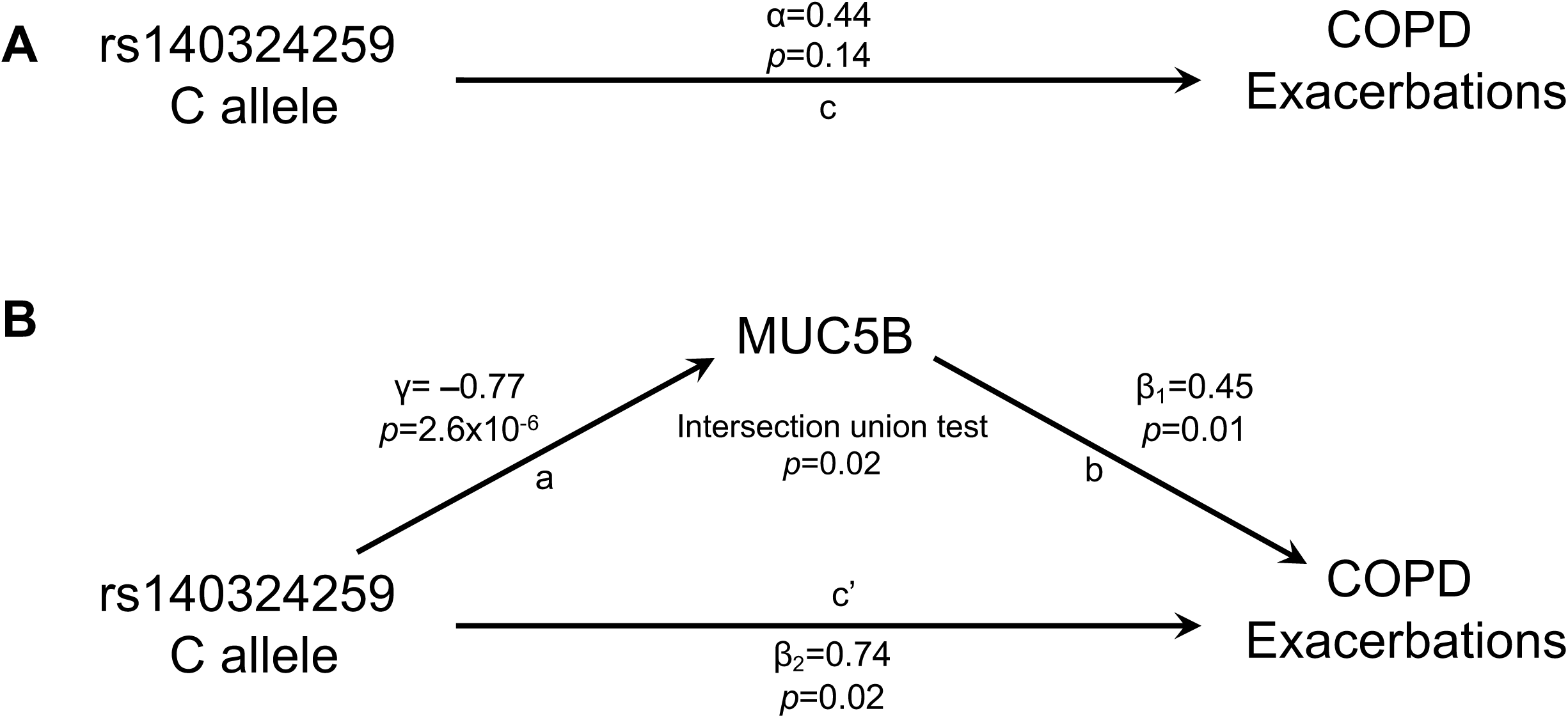
Mediation analysis reveals that rs140324259 exerts effects on exacerbations in the year prior to enrollment through direct and indirect paths with contrasting allele effects. We leveraged the mediation analysis framework of Baron and Kenny [20] to examine whether rs140324259 exerts effects on exacerbations through MUC5B. Using complete data on 142 subjects, in **(A)** we tested for the total effect of rs140324259 on acute exacerbations of COPD (“c”). In (**B)**, the mediation analysis framework is shown in which the effect of rs140324259 on acute exacerbations is modeled as the sum of direct (rs140324259 to exacerbations, c’) and indirect paths (rs140324259 to exacerbations via MUC5B (a, b)). Statistical evidence of the indirect path assessed by jointly testing that both rs140324259 → MUC5B (a) and MUC5B → exacerbations (b) are significant using an intersection union test (which is equivalent to testing that γ x β_1_ is not equal to 0). β_1_ (b) and β_2_ (c’) come from the same negative binomial regression model including both rs140324259 and MUC5B as predictors of exacerbations. Note that in this mediation analysis framework, the total effect (c) in part A is the sum of the direct (c’) and indirect paths (a→b) in part B, i.e., c = c’ + (a x b). Thus, because the sign of path a is negative while both b and c’ are positive, the total effect c (in panel A) is necessarily weaker in magnitude.

These results suggested the possibility that rs140324259 may exert effects on AE through both direct and indirect paths, the latter via MUC5B (Figure 4B). To examine this further, we employed a mediation analysis approach, based on the framework developed by Baron and Kenny [20], in which the effect of rs140324259 on AE is modeled as the sum of direct (rs140324259 → AE) and indirect paths (rs140324259 → AE via MUC5B). We leveraged an intersection union test [21] to jointly test that both components of the indirect path (rs14032425 → MUC5B and MUC5B → AE) are statistically significant. Indeed, we found evidence that this is the case (p=0.02), which is consistent with a model of partial mediation by MUC5B. Thus, overall, we conclude from these results that rs140324259 likely affects AE in two ways, both directly and indirectly, but with contrasting allele effects in each case, and overall the net effect is that rs140324259 C allele confers increased risk of AE. We note also that contrasting direction of effects of rs140324259 → MUC5B (−) and MUC5B → AE (+) likely explain why the magnitude of the association between rs140324259 and AE is weak and therefore not statistically significant (Figure 4A) [20].

We then examined associations between rs140324259 and clinical phenotypes in the larger SPIROMICS population for which genotype and clinical data exist but there is not sputum mucin concentration data (n≈1,250). In this sample, rs140324259 was associated with CB at baseline (p=0.02, Table 2). Similar to results in the smaller subset of subjects described above, the C allele was associated with increased risk of CB (odds ratio (OR) = 1.42; 95% confidence interval (CI): 1.10-1.80). The effect of rs140324259 on AE in the larger SPIROMICS sample with clinical data was examined using both retrospectively and prospectively ascertained data. rs140324259 was not significantly associated with AE in the year prior to enrollment (Table S6), nor in the year following enrollment (Table S7). However, in both of these analyses, the results were suggestive and the direction of effect was again positive for the C allele.

**Table 2.** Logistic Regression Model of Chronic Bronchitis at Baseline and rs140324259 genotype (n=1257)

| **Parameter** | **Odds Ratio** | **95% Confidence Interval** |
| --- | --- | --- |
| Age | 0.99 | 0.98-1.00 |
| Sex (M vs. F) | 1.40 | 1.09-1.80 |
| PC1 | 1.01 | 0.87-1.20 |
| PC2 | 1.10 | 0.95-1.30 |
| Smoking pack years | 1.00 | 1.00-1.00 |
| Current smoker | 5.55 | 4.17-7.40 |
| FEV1, % predicted | 0.99 | 0.99-1.00 |
| **rs140324259 (C vs. T)** | 1.42 | 1.10-1.80 |

We then asked whether rs140324259 genotype was associated with AE over a period of three years of follow up. SPIROMICS participants exacerbation frequency was categorized based on a previous study as never (n=433), inconsistent (n=331) or consistent (n=58) over the three years [22] (see Methods). Using a proportional odds model, we analyzed whether rs140324259 genotype distinguished never exacerbators versus inconsistent and consistent AE, and whether rs140324259 genotype distinguished consistent exacerbators versus never and inconsistent exacerbators. We found that rs140324259 genotype was associated with the former contrast (p=0.03), with the C allele conferring increased risk of being either an inconsistent or consistent exacerbator (Tables S8 and S9). rs140324259 genotype was not associated with the contrast between consistent exacerbators versus never and inconsistent exacerbators. To simplify the interpretation of the effect of rs140324259 on prospectively ascertained AE, we then dichotomized subjects into two groups, those who experienced AE (inconsistent and consistent) versus those that did not. As shown in Table 3, the rs140324259 C allele conferred increased risk of AE over three years of follow up (OR=1.41; 95% CI: 1.02-1.94).

**Table 3.** Logistic Regression Model Comparing Exacerbators Versus Non-Exacerbators Based on Prospectively Ascertained Exacerbation Count Over a Three-Year Period (n=822)

| **Parameter** | **Odds Ratio** | **95% Confidence Interval** |
| --- | --- | --- |
| Age | 1.00 | 0.97-1.20 |
| Sex (M vs F) | 0.65 | 0.47-0.89 |
| PC1 | 0.95 | 0.78-1.15 |
| PC2 | 1.11 | 0.90-1.37 |
| Smoking, pack years | 1.01 | 1.00-1.01 |
| Current smoker | 1.22 | 0.86-1.74 |
| Exacerbations, year prior to enrollment | 1.96 | 1.54-2.49 |
| FEV1, % predicted | 0.97 | 0.96-0.98 |
| **rs140324259 (C vs. T)** | **1.41** | **1.02-1.94** |

### Replication analyses

Finally, we analyzed data from the COPDGene study population and UK Biobank in an attempt to replicate results from SPIROMICS. For COPDGene, we utilized phenotype data from COPD cases of European ancestry, and genotype data was based on whole genome sequencing from TOPMed. Sample sizes in these analyses ranged from 5300-5700 depending on the outcome. In this population of COPD cases, rs140324259 was not associated with CB (OR = 1.08 (95% CI: 0.94-1.24), nor AE (Supplementary Tables S10-S12), though we were unable to directly evaluate whether rs140324259 was associated with the exacerbation frequency categories (never, inconsistent, consistent) described in SPIROMICS. In the UK Biobank, however, we found that rs140324259 was associated with two CB-related phenotypes, namely bringing up phlegm/sputum/mucus on most days and cough on most days (Table 4). Importantly, the C allele was enriched in cases vs. controls for these two phenotypes, thus these results are directionally consistent with results from SPIROMICS. Results for other variants in LD are shown in Table S13. As UK Biobank results were not adjusted for smoking, we additionally assessed whether rs140324259 was associated with smoking. rs140324259 was either not associated with smoking history variables or was weakly associated, but in these cases the C allele frequency was higher in controls than cases (Table 4), suggesting that the associations between rs140324259 and CB-related phenotypes in the UK Biobank are unlikely to be mediated by, or confounded with, smoking.

**Table 4.** Association analysis results for lead MUC5B pQTL variant with CB-related phenotypes and smoking history in the UK Biobank*.

| **UK Biobank Phenotype** | **# cases/controls** | **rs140324259 C**<br>**Allele Frequency**<br>**Cases** | **rs140324259 C**<br>**Allele Frequency**<br>**Controls** | **P-value** |
| --- | --- | --- | --- | --- |
| Bring up phlegm/sputum/mucus on most days | 9,250 / 97,072 | 0.150 | 0.143 | 9.80E-03 |
| Cough on most days | 14,606 / 91,635 | 0.149 | 0.143 | 4.50E-03 |
| Current Smoking Status | 43,192 / 375,625 | 0.141 | 0.143 | 9.40E-02 |
| Ever Smoked | 253,507 / 165,353 | 0.143 | 0.143 | 9.73E-01 |
| Smokes tobacco on all or most days | 2,447 / 103,281 | 0.137 | 0.144 | 1.93E-01 |

\*analysis limited to subjects of European Ancestry.

## DISCUSSION

Using quantitative measurements of sputum mucin concentrations, we identified three genome-wide significant loci and one highly suggestive locus associated with MUC5AC or MUC5B. The strongest signal we detected, with rs140324259, accounted for a large percent of variation in MUC5B, and is independent of the common *MUC5B* promoter variant associated with IPF. Surprisingly, rs140324259 does not appear to be an eQTL for *MUC5B*, though we note that our sample size for eQTL analysis was not large and that the tagSNP we used is not in very high LD with rs140324259. The mechanism by which rs140324259 (or a variant in LD) acts as a pQTL remains to be determined. One nearby variant, rs11604917, is intriguing given that it potentially disrupts binding of the transcription factor RBP-J, a key player in the Notch signaling pathway that determines ciliated vs. secretory cell fate in murine airways [19]. This could suggest that the MUC5B pQTL is a function of cell type composition of the airway epithelium, an idea supported by the lack of an association with gene expression. However, this variant is in low LD with rs140324259, and the association of rs11604917 with CB-related phenotypes in the UK Biobank was not nearly as strong as for rs140324259, arguing against a causal role for rs11604917. Still, future studies will need to address the possibility that this locus affects sputum MUC5B concentration without a corresponding effect on *MUC5B* mRNA, which has been observed for other genes [23,24].

Given that previous studies have identified eQTL for *MUC5AC* [10–12] in asthma and *MUC5B* [13] in IPF located near the genes themselves (“local eQTL”), one potential *a priori* prediction could have been that these same variants would be associated with MUC5AC and MUC5B protein concentrations. This was not the case, even in the context of a regional association analysis (i.e., not a genome-wide significance threshold). This is perhaps not surprising for at least two reasons. First, there are clear differences between our study and the previous studies as a function of disease state (COPD vs. asthma vs. IPF) and anatomical location (upper vs. lower airways). Second, mucin protein concentration is the product of several pathways beyond just mucin gene transcription, including protein synthesis, post-translation modifications, packaging into vesicles, secretion, airway hydration via ion transport, and mucociliary clearance. Thus, one could reasonably expect that genetic variants that regulate any of these processes could be associated with mucin concentration. It remains to be determined whether any of the distal/off-chromosome pQTL identified here play a role in one or more of these pathways. That we did not identify any associations in/near genes with known roles in these processes suggests that either we were underpowered to detect these associations and/or that there is limited functional genetic variation in/near these genes.

Our analysis of associations between rs140324259 and clinical outcomes, namely CB and AE, produced intriguing results in the SPIROMICS cohort, including that the variant was associated with AE ascertained prospectively over three years. These results did not replicate in COPDGene, but we did find an association with sputum production and cough in a much larger dataset, the UK Biobank. These data argue in support of a role for the MUC5B pQTL in CB-related phenotypes. However, we acknowledge that the results of CB and AE association analyses with rs140324259 in SPIROMICS would not a survive multiple testing correction based on the number of outcomes/models we evaluated; in addition, we were unable to replicate these results COPDGene, thus raising the potential that the results in SPIROMICS represent false positives. We note here that failure to replicate genetic associations with AE is unfortunately common [25], and future studies in which standardized definitions of AE can be employed will certainly facilitate the best comparisons across studies [25]. It is also worth noting that a previous study identified significant blood biomarkers of susceptibility for AE in SPIROMICS and separately in COPDGene, but there was essentially no overlap in associations between the two populations [26], which points to the difficulty in identifying reproducible predictors of exacerbations. The UK Biobank analysis, while supportive of our results, did not have the same degree of detailed respiratory phenotypes and was performed in a general population sample.

Additional analyses adjusting for disease state and other covariates could be beneficial [27]). Further attempts to replicate these finding in other populations would also be useful, in particular to address the question of generalizability across populations of different genetic ancestries.

In aggregate, the results of association tests between rs140324259 and COPD phenotypes suggest an apparent paradox. While MUC5B concentration was positively associated with AE and CB in SPIROMICS, and the C allele was associated with significantly reduced MUC5B, the C allele overall was associated increased risk of AE and CB. This result suggests that higher expression of MUC5B may in fact be protective against AE and CB, perhaps by virtue of normalizing the ratio of elevated MUC5AC to MUC5B, making it more clearable, as has been suggested before in relationship to the IPF-associated variant rs35705950 [28].

While we examined associations between loci associated with mucins, CB, and AE in COPD patients specifically, others have examined the genetics of CB/chronic mucin hypersecretion in combined analysis of the general population and patients with COPD [27,29] or in smokers without COPD [30]. In the study with COPD cases and the general population [29], the most consistent association signal was for rs6577641, which was also shown to act as an eQTL for the gene *SATB1*. In look up analysis, this variant was not associated with either sputum MUC5B concentration or CB in the SPIROMICS population, nor was the lead variant (rs10461985) from another study [30]. The most recent study reported an association of variants on proximal Chr 11 (near *MUC2*) with chronic sputum production using the same UK Biobank phenotype codes we used [27], but the LD between lead variant in that study (rs779167905) and rs140324259 is minimal (R^2^=0.08), making it unlikely that these are the same signals.

In summary, we identified pQTL for MUC5AC and MUC5B in sputum, demonstrating that common genetic variants influence these biomarkers. The lead MUC5B pQTL, rs140324259, was associated with CB and prospectively ascertained AE in SPIROMICS and was also associated with CB-related phenotypes in the UK Biobank. Additional studies are needed to further evaluate whether rs140324259 may be a biomarker of CB and AE susceptibility in COPD in other populations, and to determine how this variant influences MUC5B concentration in sputum.

## MATERIALS AND METHODS

### Ethics statement

Subjects provided informed consent to participate in the studies described here. Details and institutional review boards for each clinical site are provided in Supporting File 1.

### Study subjects and genotype data

The primary analyses presented here are based on study participants in SPIROMICS (ClinicalTrials.gov Identifier: NCT01969344), and a schematic of the SPIROMICS datasets used here is shown in Figure S1. The study design has been described previously [31]. SPIROMICS participants were genotyped using the Illumina OmniExpress Human Exome Beadchip [32]. Quality controls included testing for sex concordance and removal of SNPs with high genotype missing rates (>5%) and/or Hardy Weinberg p < 1×10^−6^. Genotype imputation was performed using the Michigan Imputation Server [33] using haplotypes from Phase3 of the 1000 Genomes Project [34]. Study participants were categorized into either European ancestry (EA, N=576 or African ancestry (AA, N=132) groups based on genotype data. We used an adaptive R^2^ threshold to filter imputed variants in each ancestry group based on the minor allele frequency (MAF). For each MAF interval, the R^2^ value was chosen such that the average R^2^ for variants with values larger than the threshold is at least 0.8 (Tables S14 and S15). We limited our analyses to SNPs with minor allele counts >8, resulting in ~10 million SNPs in EA and 12 million variants in AA subjects for association with total and specific mucin concentrations.

In COPDGene [35] (ClinicalTrials.gov Identifier: NCT00608764), genotype data for rs140324259 was obtained from whole genome sequencing performed through the TOPMed consortium [36]. Results from the UK Biobank data were obtained from the Pan-UK Biobank analysis (see further description below) [37].

### Sputum Mucin Phenotype data

Sputum mucin concentration: sputum induction and measurement methods have been previously reported [8,9,38]. In brief, hypertonic saline was used to induce sputum, which was then placed in a buffer containing 6 molar guanidine, and stored at 4 degrees. Total sputum mucin concentration was determined using a size exclusion chromatography / differential refractometry measurement approach. For a subset of subjects, MUC5AC and MUC5B concentration was determined using stable isotope labeled mass spectrometry [38]. Data were generated in two batches. In addition to SPIROMICS participants with COPD (n=439), two additional sets of subjects were also included in the mucin analyses: non-smoking controls (n=50), and smokers without COPD that are referred to as the “at-risk” group (n=219). These subjects were included in genetic analysis of sputum mucin concentration but were not included in the analysis of COPD outcomes.

### Clinical/Phenotype Data

We analyzed data on two COPD phenotypes, namely CB and AE, in SPIROMICS. The CB phenotype was ascertained at the first study visit (“baseline”) and was categorized based on participants’ responses to questions regarding frequency of cough and mucus/phlegm production in the St. George’s Respiratory Questionnaire. The analysis of AE was based on previous work from SPIROMICS [22,31], in which AE were defined as events that required health care utilization (i.e., office visit, hospital admission, or emergency department visit for a respiratory flare-up) involving the use of antibiotics or systemic corticosteroids, or both. In COPDGene, chronic bronchitis was based on chronic cough and phlegm production for ≥ 3 mo/y for 2 consecutive years [5]. For AE, self-reported moderate-to-severe exacerbations in the year prior to enrollment and the number of moderate-to-severe exacerbations ascertained prospectively from longitudinal follow up data were examined. In the Pan-UK Biobank (https://pan.ukbb.broadinstitute.org/), we evaluated results of association analyses for two CB-related phenotypes, namely bringing up phlegm/sputum/mucus daily (yes vs. no, phenocode 22504) and coughing on most days (yes vs. no, phenocode 22502), as well as smoking history variables, of which were assessed by questionnaire.

### Statistical models

#### Sputum mucin concentration

Data on total and specific (MUC5AC and MUC5B) mucin concentrations were log-transformed prior to analysis. GWAS analysis was performed using version 0.5.0 of the ProbABEL software [39]. Analysis of total mucin concentration was conducted in each ancestry group separately (N=576 EA and 132 AA), followed by a pooled analysis of both ancestry groups. In ancestry-specific analyses, main effect SNP models of each mucin phenotype included covariates for the top two principal components of ancestry (PC) obtained from EIGENSTRAT [40], age, sex, batch of mucin quantitation analysis, current smoking status, smoking pack-years, and CB. Results were not materially different when we included up to 10 genotype PCs. For MUC5AC and MUC5B, GWAS was performed in EA subjects only (N=215) with the same covariates used for total mucin concentration. SNP-based heritability for MUC5AC and MUC5B was estimated on an LD-pruned set of markers from the genotyped data (subsetting to individuals with the relevant phenotype data) using GCTA version 1.92.1. We performed exploratory genome-wide interaction studies of SNP × smoking interactions in which we tested for the joint effects of SNP and SNP × smoking interactions (2 d.f. test) on mucin concentrations in models including the same covariates as above.

#### eQTL Analysis

Airway epithelia gene expression from 144 SPIROMICS participants was analyzed to test whether rs140324259 is an eQTL for *MUC5B* by performing a genome-wide eQTL mapping as described before in Kasela et al. [15]. Briefly, RNA-seq data from the airway epithelium was normalized, filtered, and transformed using inverse normal transformation.

Genotype data was obtained from TOPMed (Freeze 9) [36]. The eQTL regression model for a given gene included sex, four genotype PCs, and 15 PEER factors (probabilistic estimation of expression residuals [41]) as covariates. eQTL mapping was performed using tensorQTL [42] and 10,000 permutations were used to control for multiple testing at false discovery rate (FDR) < 0.05. To look up the eQTL association with *MUC5B*, we used the proxy SNP rs55680540 because rs140324259 did not pass variant filter quality control.

#### Clinical Phenotypes

Chronic Bronchitis (CB): Following the analyses of sputum mucin concentration data, we tested for an association between the lead variant for sputum MUC5B concentration (rs140324259) and CB in the larger SPIROMICS population (N=1257). Logistic regression models were used for CB, accounting for the top two genotype PCs, age, sex, current smoking status, pack-years of smoking, and FEV1.

Acute Exacerbations (AE): Exacerbation outcomes were modeled using negative binomial regression models including the same covariates as above. Additionally, in the analysis of prospectively ascertained AE, we included AE in the year prior to enrollment as a predictor. Because prior work showed that exacerbation frequency among subjects with COPD in SPIROMICS is not stable [22], we leveraged a previously developed classification system which categorized SPIROMICS participants as never, inconsistent or consistent exacerbators using three years of follow up data [22]. Consistent exacerbators were subjects who experienced at least one acute exacerbation in each of the three years; subjects who had an exacerbation during some but not all of the three years of follow up where defined as inconsistent exacerbators. We analyzed the association between rs140324259 genotype and these three exacerbation groups using a proportional odds model, comparing (1) never exacerbators versus inconsistent and consistent exacerbators, and (2) consistent exacerbators versus never and inconsistent exacerbators. Based on the results of these analyses, we collapsed the exacerbation groups into two categories: ever (combining inconsistent and consistent exacerbators) vs. never exacerbators, then modeled this outcome using logistic regression with covariates for top two genotype PCs, age, sex, current smoking status, pack-years of smoking, FEV1 (% predicted), and the number of AE in the year prior to enrollment.

#### Mediation analysis

in the subset of SPIROMICS subjects for which there is complete phenotype data on genotype, sputum MUC5B, and clinical outcomes (N=141), we tested for evidence of that MUC5B mediates an association between rs140324259 and AE (i.e. rs140324259 → MUC5B → AE), invoking the overall mediation analysis framework of Baron and Kenny [20].

We evaluated a direct path from rs140324259 → AE (c), and an indirect path from rs140324259 → MUC5B (a) and MUC5B → AE (b), while also examining the path from rs140324259 → AE conditional on MUC5B (c’). All regression models included age, sex, two ancestry PCs, current smoking status, pack-years of smoking, and FEV1 (% predicted) as predictors. For (a), we used a linear model for MUC5B in which rs140324259 was coded linearly (0,1,2). For (b), (c), and (c’), we used we used negative binomial regression models of AE that included rs140324259 (c), MUC5B (b), or both (c’). To formally test for mediation, we leveraged the SNP mediation intersection-union test (SMUT) [21] which jointly tests for non-zero parameter estimates from models for (a) and (b), which is equivalent to testing that a x b is not equal to 0 in the Baron and Kenny framework.

## Acknowledgements

Y.L. was partially supported by NIH grants R01GM105765 and U01DA052713. S.K. and T.L. were supported by NIH grant R01HL142028. S.N.P.K. was partially supported by NIH grant R01 HL122711.

The authors thank Dr. Wesley Crouse for helpful discussions related to mediation analysis. The authors are grateful to SPIROMICS participants and participating physicians, investigators and staff, for making this research possible. More information about the study and how to access SPIROMICS data is available at www.spiromics.org. The authors would like to acknowledge the University of North Carolina at Chapel Hill BioSpecimen Processing Facility for sample processing, storage, and sample disbursements (http://bsp.web.unc.edu/).

We would like to acknowledge the following current and former investigators of the SPIROMICS sites and reading centers: Neil E Alexis, MD; Wayne H Anderson, PhD; Mehrdad Arjomandi, MD; Igor Barjaktarevic, MD, PhD; R Graham Barr, MD, DrPH; Patricia Basta, PhD; Lori A Bateman, MSc; Surya P Bhatt, MD; Eugene R Bleecker, MD; Richard C Boucher, MD; Russell P Bowler, MD, PhD; Stephanie A Christenson, MD; Alejandro P Comellas, MD; Christopher B Cooper, MD, PhD; David J Couper, PhD; Gerard J Criner, MD; Ronald G Crystal, MD; Jeffrey L Curtis, MD; Claire M Doerschuk, MD; Mark T Dransfield, MD; Brad Drummond, MD; Christine M Freeman, PhD; Craig Galban, PhD; MeiLan K Han, MD, MS; Nadia N Hansel, MD, MPH; Annette T Hastie, PhD; Eric A Hoffman, PhD; Yvonne Huang, MD; Robert J Kaner, MD; Richard E Kanner, MD; Eric C Kleerup, MD; Jerry A Krishnan, MD, PhD; Lisa M LaVange, PhD; Stephen C Lazarus, MD; Fernando J Martinez, MD, MS; Deborah A Meyers, PhD; Wendy C Moore, MD; John D Newell Jr, MD; Robert Paine, III, MD; Laura Paulin, MD, MHS; Stephen P Peters, MD, PhD; Cheryl Pirozzi, MD; Nirupama Putcha, MD, MHS; Elizabeth C Oelsner, MD, MPH; Wanda K O’Neal, PhD; Victor E Ortega, MD, PhD; Sanjeev Raman, MBBS, MD; Stephen I. Rennard, MD; Donald P Tashkin, MD; J Michael Wells, MD; Robert A Wise, MD; and Prescott G Woodruff, MD, MPH. The project officers from the Lung Division of the National Heart, Lung, and Blood Institute were Lisa Postow, PhD, and Lisa Viviano, BSN; SPIROMICS was supported by contracts from the NIH/NHLBI (HHSN268200900013C, HHSN268200900014C, HHSN268200900015C, HHSN268200900016C, HHSN268200900017C, HHSN268200900018C, HHSN268200900019C, HHSN268200900020C), grants from the NIH/NHLBI (U01 HL137880 and U24 HL141762), and supplemented by contributions made through the Foundation for the NIH and the COPD Foundation from AstraZeneca/MedImmune; Bayer; Bellerophon Therapeutics; Boehringer-Ingelheim Pharmaceuticals, Inc.; Chiesi Farmaceutici S.p.A.; Forest Research Institute, Inc.; GlaxoSmithKline; Grifols Therapeutics, Inc.; Ikaria, Inc.; Novartis Pharmaceuticals Corporation; Nycomed GmbH; ProterixBio; Regeneron Pharmaceuticals, Inc.; Sanofi; Sunovion; Takeda Pharmaceutical Company; and Theravance Biopharma and Mylan.

The COPDGene project described was supported by Award Number U01 HL089897 and Award Number U01 HL089856 from the National Heart, Lung, and Blood Institute. The content is solely the responsibility of the authors and does not necessarily represent the official views of the National Heart, Lung, and Blood Institute or the National Institutes of Health. The COPDGene project is also supported by the COPD Foundation through contributions made to an Industry Advisory Board comprised of AstraZeneca, Boehringer Ingelheim, GlaxoSmithKline, Novartis, Pfizer, Siemens and Sunovion. A full listing of COPDGene investigators can be found at: http://www.copdgene.org/directory

Molecular data for the Trans-Omics in Precision Medicine (TOPMed) program was supported by the National Heart, Lung and Blood Institute (NHLBI). Genome sequencing for “NHLBI TOPMed: Genetic Epidemiology of COPD (COPDGene)” (phs000951.v4.p4) was performed at the Northwest Genomics Center (3R01HL089856-08S1) and Broad Institute Genomics Platform (HHSN268201500014C). Core support including centralized genomic read mapping and genotype calling, along with variant quality metrics and filtering were provided by the TOPMed Informatics Research Center (3R01HL-117626-02S1; contract HHSN268201800002I). Core support including phenotype harmonization, data management, sample-identity QC, and general program coordination were provided by the TOPMed Data Coordinating Center (R01HL-120393; U01HL-120393; contract HHSN268201800001I). We gratefully acknowledge the studies and participants who provided biological samples and data for TOPMed.

