## Supplemental Figures for "Genetic Regulators of Sputum Mucin Concentration and Their Associations with COPD Phenotypes"

#### Figure S1

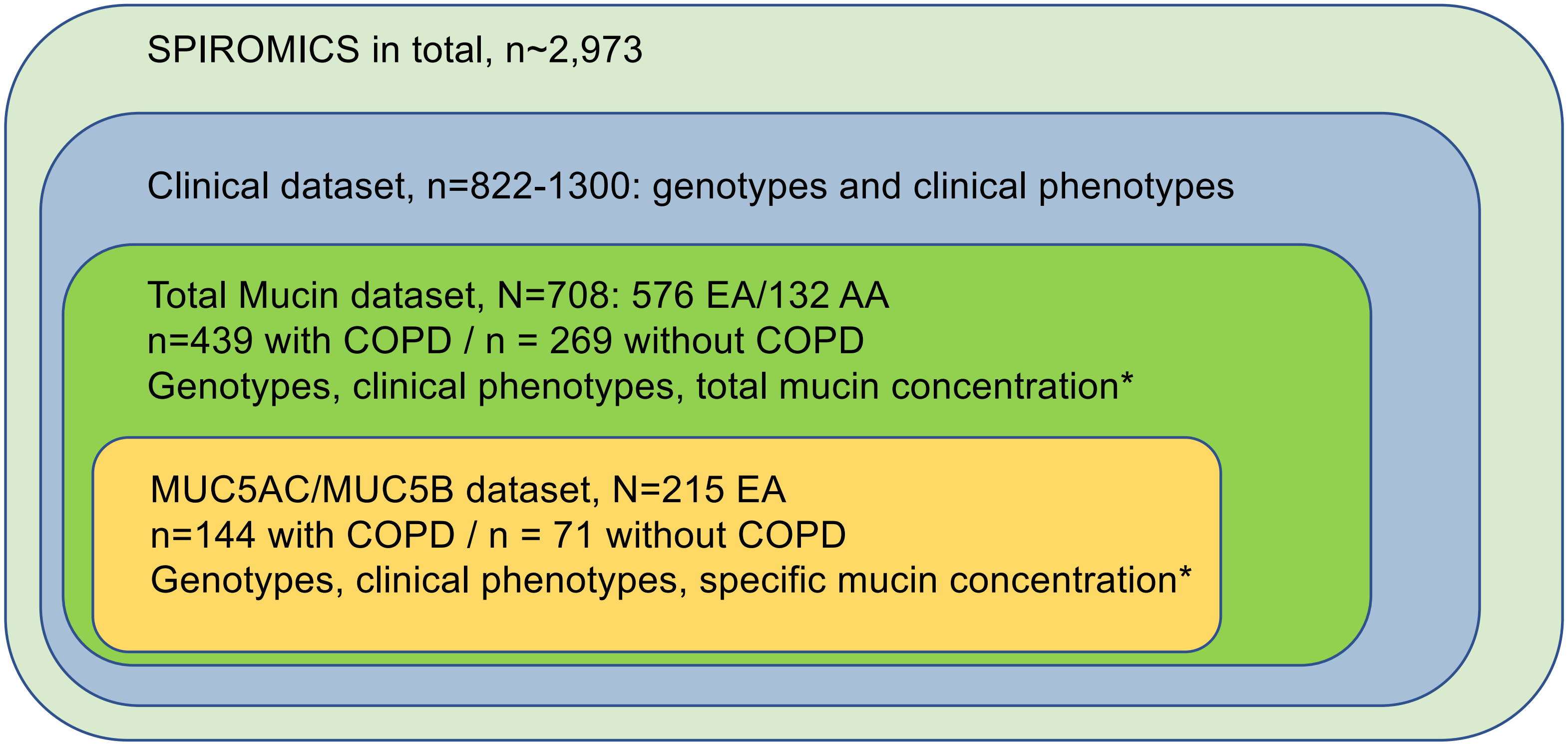


**Figure S1. SPIROMICS Participant Data Utilized in This Study.** Clinical data used includes FEV1, chronic bronchitis, and acute exacerbations. In GWAS models, all subjects were used; in models of clinical outcomes, subjects without COPD were removed. AA: African Ancestry, EA: European Ancestry. Note that while the figure suggests nested subsets of SPIROMICS data used in these analyses, in reality, there are varying degrees of overlap between subjects in mucin datasets and clinical dataset.

### Figure S2

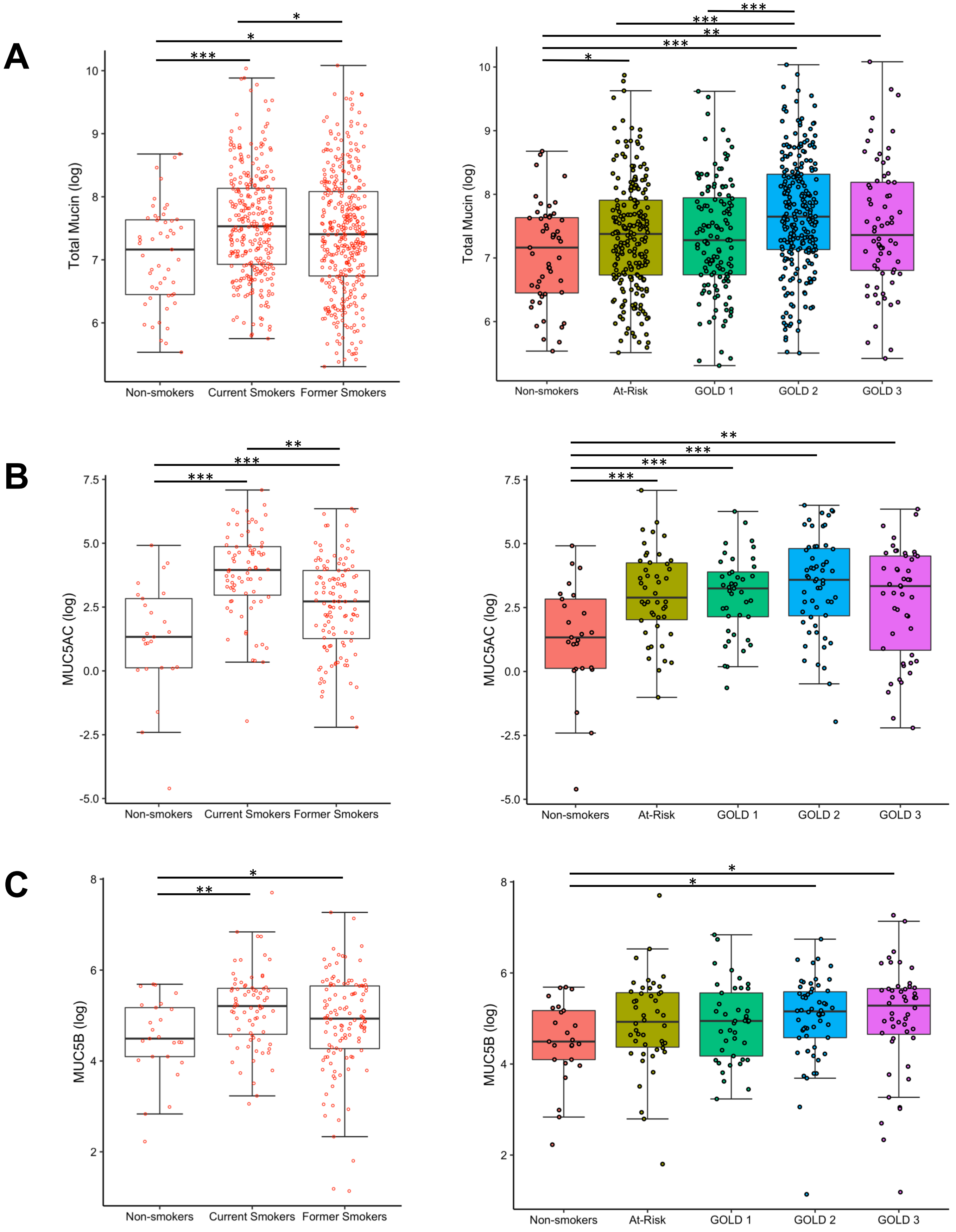


**Figure S2. Distributions of Sputum Mucin Concentrations.** Distributions for total mucin (A, n=576 EA/132 AA), MUC5AC (B, n=215 EA), and MUC5B (C, n=215) as a function of smoking history (left) and GOLD stage (right).

### Figure S3

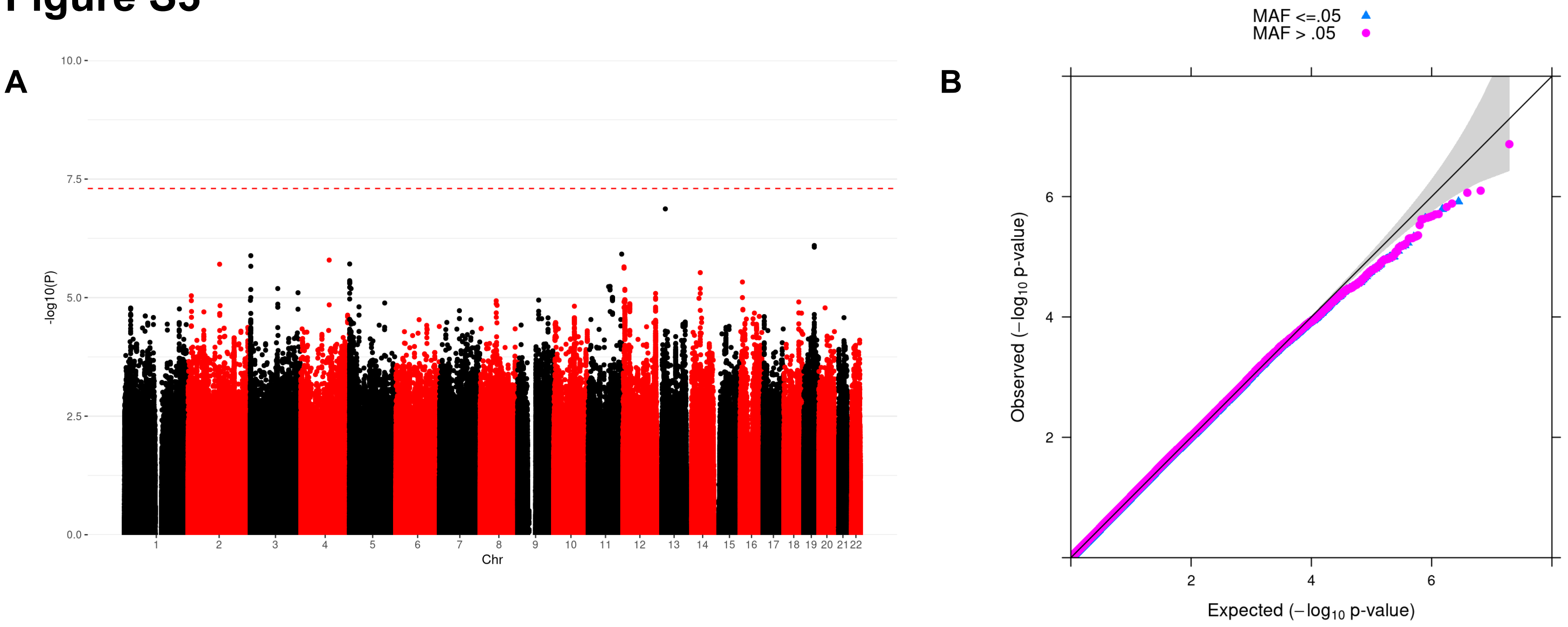


**Figure S3. GWAS results for sputum total mucin concentration in EA subjects (N=576). A**. Manhattan plot. N=576. **B**. Corresponding quantile-quantile plot.

### Figure S4

A

B

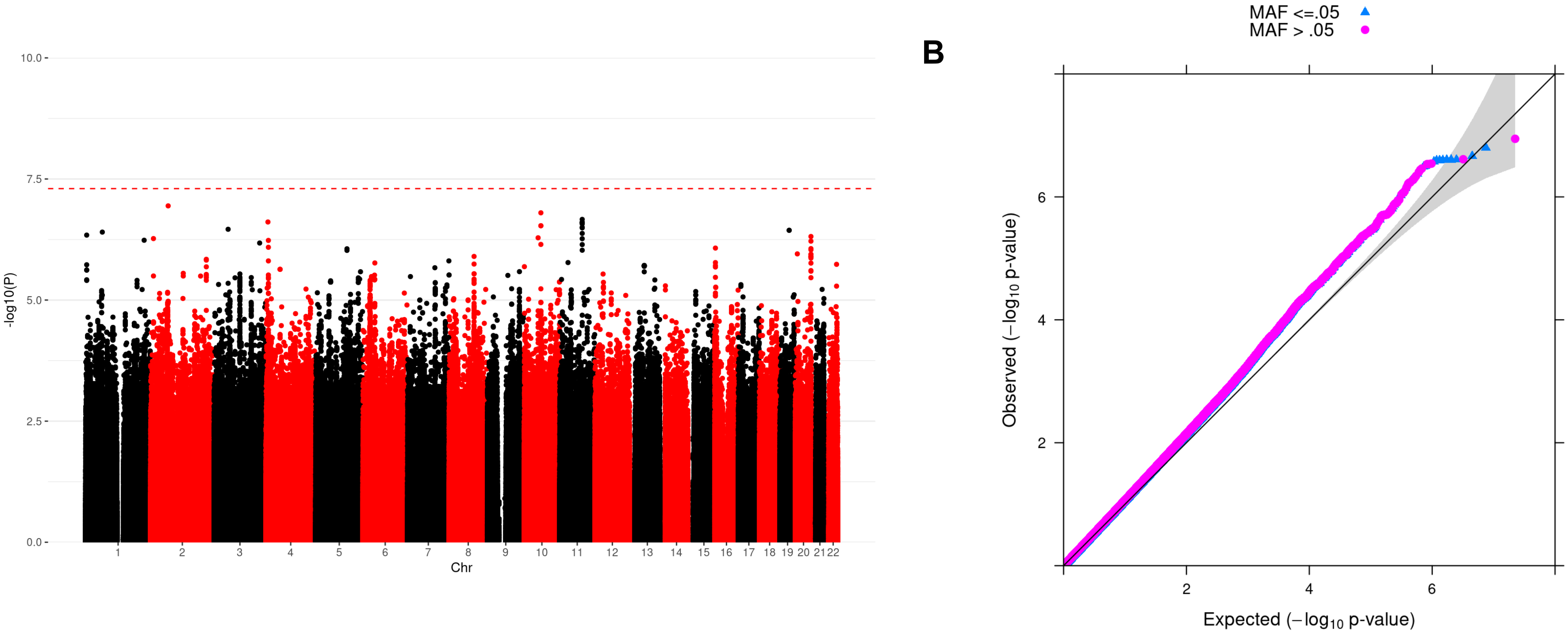


**Figure S4. GWAS results for sputum total mucin concentration in AA subjects (N=132). A**. Manhattan plot. N=132. **B.** Corresponding quantile-quantile plot.

### Figure S5

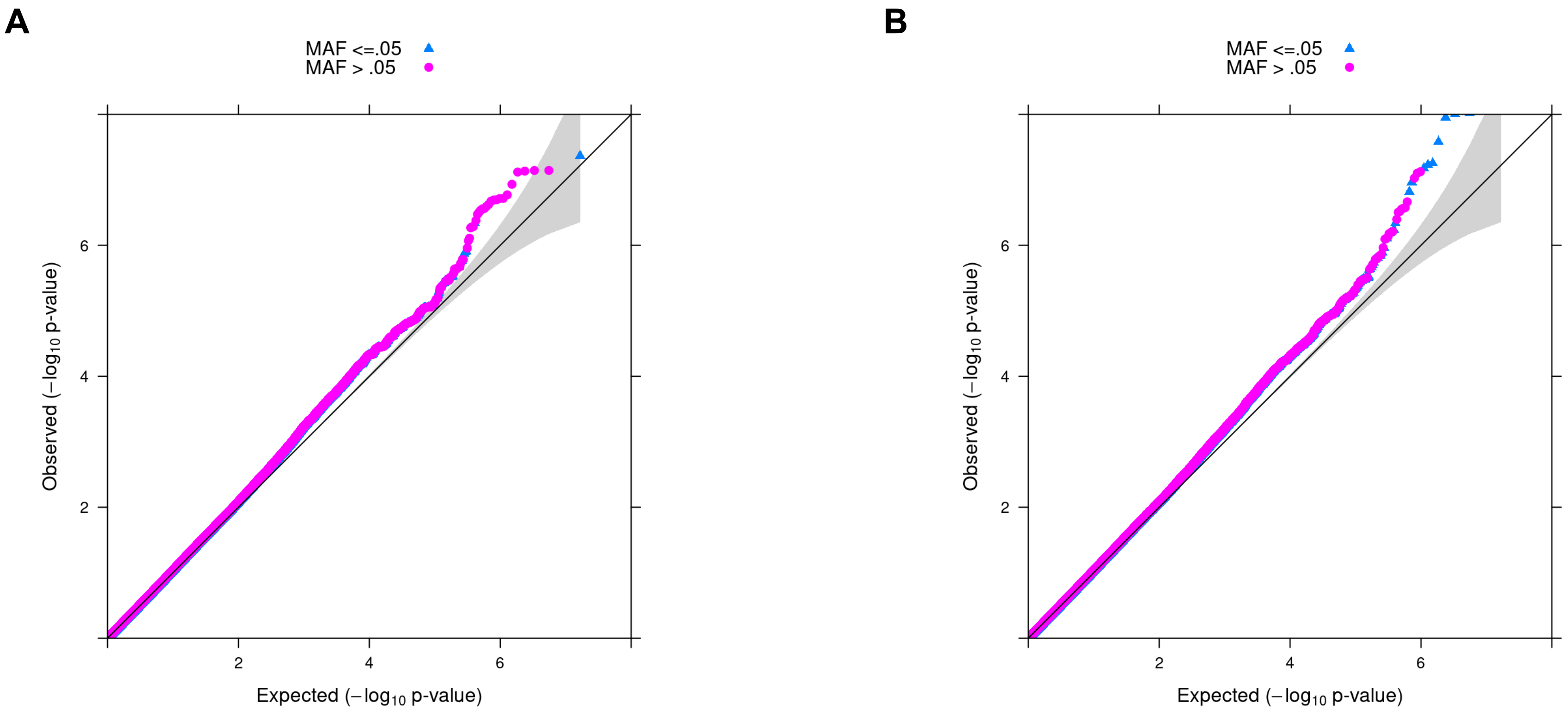


**Figure S5. Quantile-quantile plots for MUC5AC (A) and MUC5B (B) GWAS results in EA subjects (n=215).**

### Figure S6

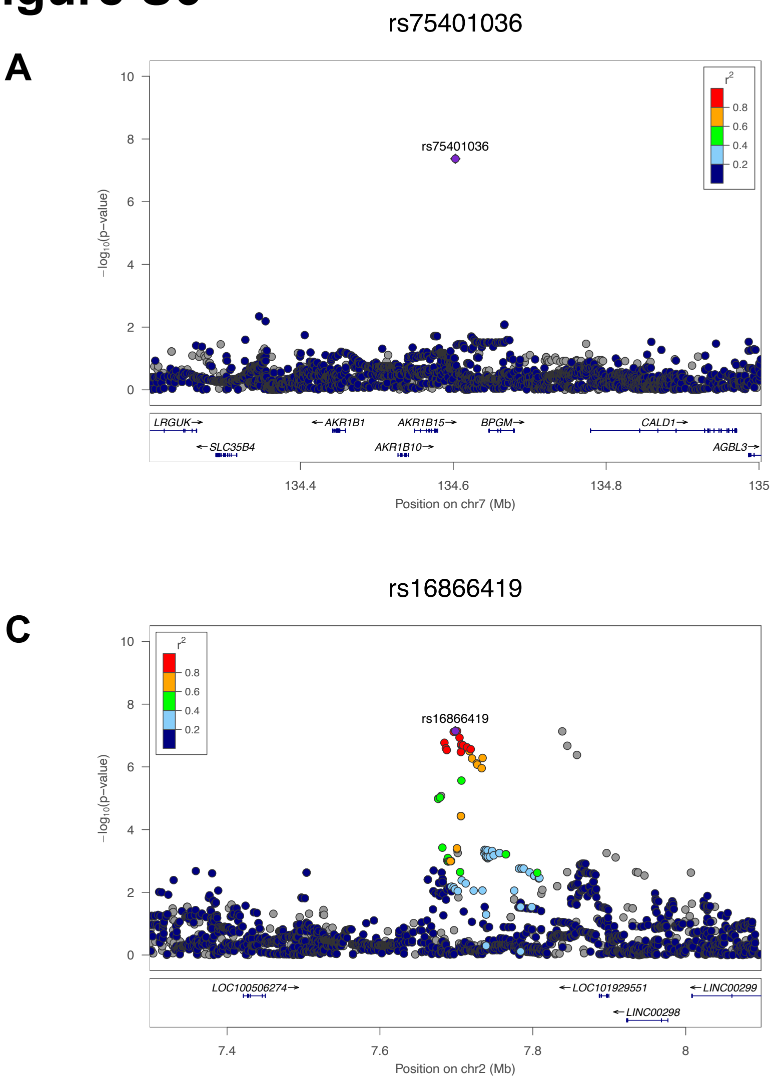


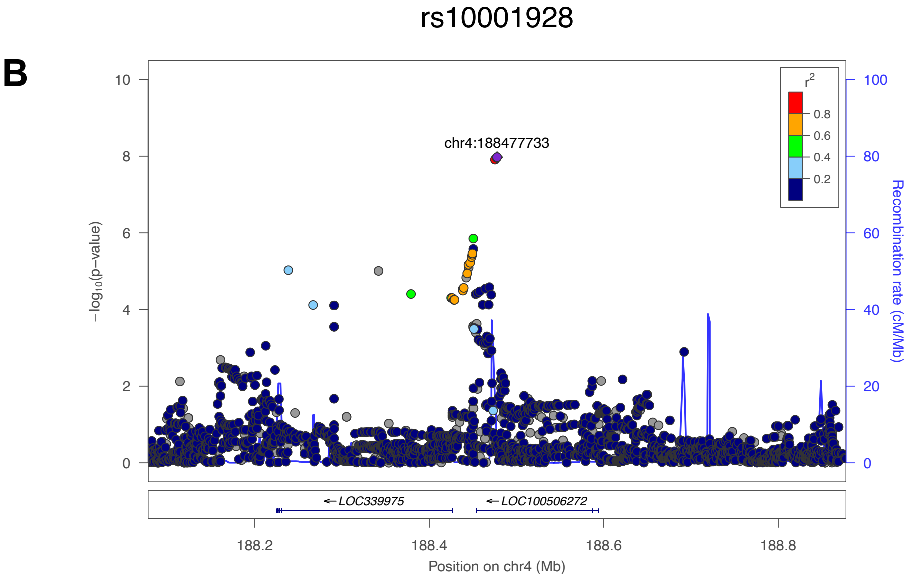


**Figure S6. Locus zoom plots for mucin pQTL that act in trans.** MUC5AC pQTL on chromosome 7 (A) and MUC5B pQTL on chromosome 4 (B). The suggestive locus for MUC5AC on Chromosome 2 is also shown (C).

### Figure S7

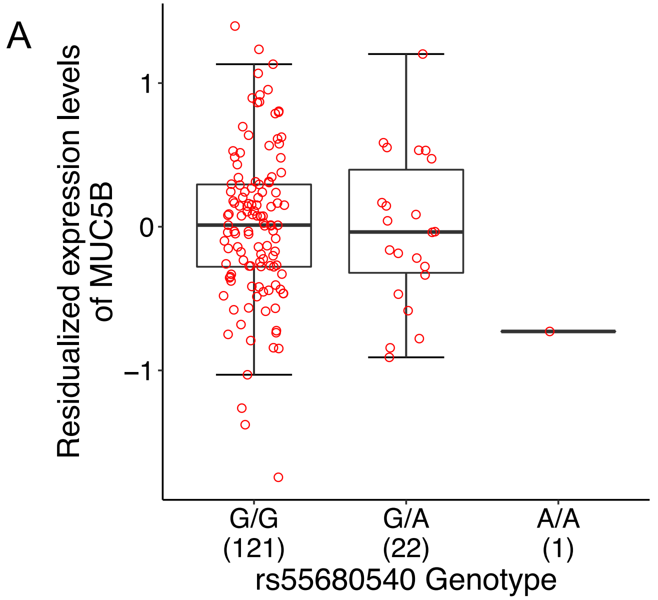


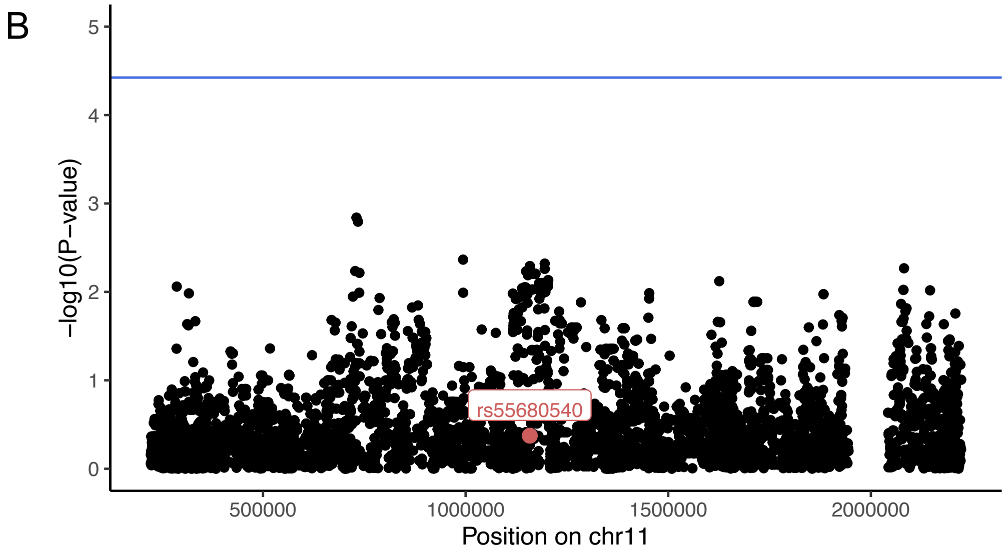


**Figure S7. eQTL analysis for *MUC5B* in 144 SPIROMICS participants. A.** Airway brush RNA-seq data was analyzed in relation to a proxy SNP for rs140324259, rs55680540. MUC5B expression data is plotted as residuals from a model containing four genotype PCs, age, sex, and 15 PEER factors. **B.** Results of scan for other local eQTL for MUC5B, with rs55680540 highlighted. Blue horizontal line corresponds to regional multi-testing threshold.

### Figure S8

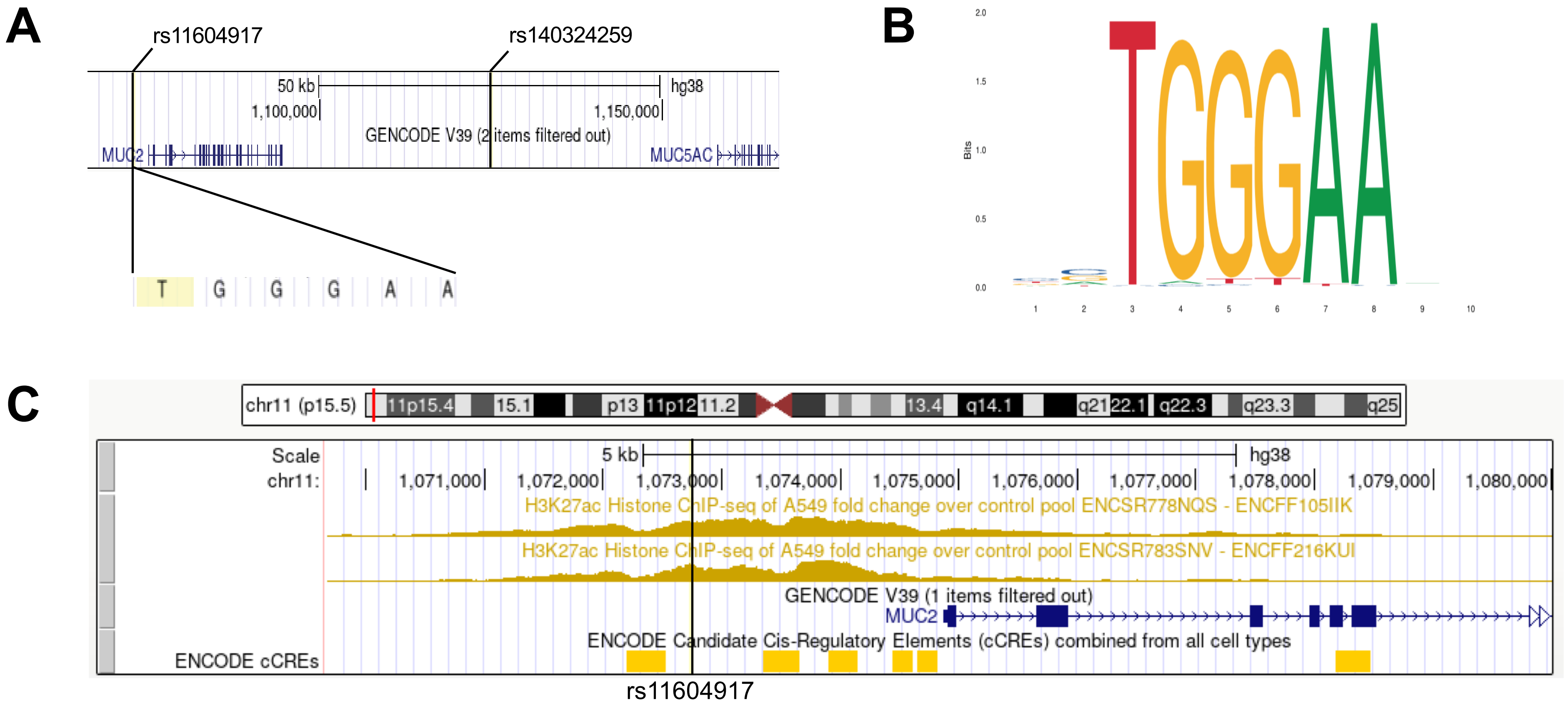


**Figure S8. Putative disruption of RBP-J binding to a region upstream of mucin gene cluster on Chr 11. A**. Location of lead MUC5B pQTL variant, rs140324259, and rs11604917. Note that rs11604917 (T->C) lies in the first position of a putative RBP-J binding motif. **B.** Consensus motif for RBP-J. Note that the first position is essentially invariant. **C.** Location of rs11604917 relative to enhancer marks detected in the airway epithelia cell line A549.
